# Scalable Isolation of Surface-Engineered Extracellular Vesicles and Separation of Free Proteins via Tangential Flow Filtration and Size Exclusion Chromatography (TFF-SEC)

**DOI:** 10.1101/2024.03.07.584007

**Authors:** Yuki Kawai-Harada, Vasudha Nimmagadda, Masako Harada

## Abstract

**Background:** Extracellular vesicles (EVs) represent small lipid bilayer structures pivotal in mediating intercellular communication via biomolecular transfer. Their inherent characteristics, including packaging, non-immunogenicity, and biofluid stability, position EVs as promising drug delivery vectors. However, developing clinical quality EVs requires multifaceted technological advancement.

**Methods:** In this study, a method is introduced for engineering extracellular vesicles (eEVs) from cultured cells and their subsequent isolation using lab-scale tangential flow filtration (TFF). This is the first study to evaluate DNA loading efficacy into EVs isolated by TFF, marking a significant milestone in the field of targeted drug delivery. Initially, cells are transfected with EV-display constructs to facilitate the secretion of eEVs bearing the desired coding molecules. Following brief centrifugation, the cell culture media undergoes filtration using hollow fiber filters. TFF, by applying a constant flow, effectively segregates molecules based on designated molecular weight cut-off (MWCO), enriching particles between 50 nm and 650 nm.

**Results:** Compared to conventional methods like ultracentrifugation, TFF demonstrates higher efficiency in removing undesired molecules/aggregates while exerting less stress on EVs. Characterization of eEVs through various assays confirms TFF’s superiority in isolating pure EV populations. Additionally, the necessity of size-exclusion chromatography (SEC) after tangential flow filtration (TFF) becomes evident for effectively removing unbound protein contaminants.

**Conclusion:** In conclusion, TFF-SEC emerges as a scalable and superior approach for eEV isolation, promising significant advancements in clinical applications.

## Introduction

Extracellular vesicles (EVs) represent heterogeneous populations of small, membrane-enclosed entities that are released into the extracellular milieu.[1] These EVs play a vital role in facilitating intracellular communication by serving as vehicles for transport of variety of molecular constituents, thereby influencing diverse pathological and physiological processes.[1-4] This cargo includes nucleic acids, proteins, lipids, and membrane receptors.[1] It is noteworthy that each EV is unique, with its composition varying based on several factors including source cell type, size, biogenesis pathway, environment, and stimuli.[3] These factors not only aid in the classification of EVs but also open avenues for engineering capabilities through their manipulation, such as altering the EV source cell.[2, 5] Additionally, it is essential to acknowledge that EV circulation is an innate biological process.[6] EVs are ubiquitously present and secreted across a broad spectrum of biological sources, including animals, plants, bacteria, and fungi.[7-9] Consequently, these vesicles are capable of efficiently ferrying molecular cargo and exerting influence on recipient cells in manners that are often unattainable through other mechanisms.[6, 10] The combination of these inherent characteristics with the potential for engineering enhancements underscores the significant promise of EVs as therapeutic delivery systems and disease biomarkers in clinical applications.[2, 6, 11]

Despite the considerable potential of EVs, the process of EV isolation posed numerous challenges and remains an area of active research interest.[6, 12-15] The objective of EV isolation is to establish a reproducible methodology for generating pure, intact EVs on a large scale.[12] While various methods have been employed for EV isolation, including ultracentrifugation (UC), size-exclusion chromatography (SEC), and ultrafiltration (UF), UC has emerged as the most widely used for isolating EVs from cell culture media. [12, 16-19] However, a universally accepted method is yet to be established, primarily due to the inherent limitations associated with each approach.[20] For example, UC, despite its widespread use, is characterized by low EV yield and purity of EVs and is time-consuming.[12] UF, while less time-intensive than UC, is prone to issues such as clogging of EVs leading to filter plugging.[21] In response to these challenges, various isolation methods are increasingly used in combination, employing a range of adapted protocols in an attempt to circumvent these limitations.[12, 22, 23]

Tangential flow filtration (TFF) is an advanced form of UF, utilizing ‘tangential flow’ where the media flows parallel to the membrane, in contrast to the perpendicular flow in UF’s dead-end filtration.[24] TFF offers advantages over UC in terms of time efficiency, scalability and need for expensive equipment.[10] Several studies have reported a scalable method for purifying EVs with low protein contamination by combining TFF and SEC based on these characteristics. However, this is the first study to both demonstrate the EV isolation method and compare DNA cargo loading.[25-27]

This study is designed to investigate the efficiency of UC and TFF methods, in conjunction with SEC, for the isolation of surface-engineered EVs as molecular transporters.

## Material and Methods

### Cell Culture and Treatment

Before the experiments, the following cell line from American Type Culture Collection (ATCC), was tested for mycoplasma: HEK293T (Human Embryonic Kidney cell line). The cells were cultured in high-glucose DMEM (Gibco) supplemented with 100U/mL penicillin, 100 μg/mL streptomycin, and 10% (v/v) fetal bovine serum (FBS, Gibco). All cells were maintained in a humidified incubator with 5% CO_2_ at 37⍰:°C. For EV production we employed the method previously described by our group with slight modifications.[28] Briefly, HEK293T cells were initially seeded at 1×10^6^ in a 10 cm tissue culture dish and allowed to grow for 24 h. The following day, a mixture containing 10 μg pDNA (pcS-RDG-C1C2, Addgene #200163) and PEI in a ratio of 1:2.5 (DNA/PEI) was prepared in non-supplemented DMEM. The mixture was then added to the cells after 30 s-pulse-vortexing and incubated for 10 min at room temperature.1 After 24 h of further incubation, the cells were washed with PBS, and the culture media was replaced with 20 mL of DMEM supplemented with Insulin-Transferrin-Selenium (ITS) (Corning), along with 100U/mL penicillin and 100 μg/mL streptomycin (conditioned media). The cells were then allowed to grow for another 24 h for engineered EV (eEV) generation. To generate Naïve EVs, HEK293T cells were seeded under the same conditions. After 48 h of further incubation, the culture medium was replaced with 20 mL of DMEM supplemented with Insulin-Transferrin-Selenium (ITS) (Corning), along with 100U/mL penicillin and 100 μg/mL streptomycin (conditioned media), and allowed to grow for an another 24 h.

### EV isolation

#### Centrifugation

For EV centrifugation, we adopted the method previously described by our group with slight modifications.[29] Briefly, EVs were purified from conditioned media by differential centrifugation at 600g for 30 min, the supernatant was further centrifuged at 2000g for 30 min, and the supernatant was further ultracentrifuged in PET Thin-Walled ultracentrifuge tubes (Thermo Scientific 75000471) at 12,000g with a Sorvall WX+ Ultracentrifuge equipped with an AH-629 rotor (k factor = 242.0) for 90 min at 4 °C to pellet the large EVs (lEVs) and at 100,000g for 90 min at 4 °C to pellet the small EVs (sEVs). The pellet containing EVs was resuspended in EV storage buffer (0.2% Bovine Serum Albumin (Life Technologies, Carlsbad, CA, USA), 25 mM D-(+)-Trehalose dihydrate (TCI America, Portland, OR, USA), and 25 mM HEPES pH7.0 in PBS).[28]

#### Tangential Flow Filtration

EVs were purified from conditioned media by differential centrifugation. Briefly, the media was centrifuged at 600g for 30 min to remove the cell and cell debris, and the supernatant was further centrifuged at 2000g for 30 min to remove apoptotic bodies. Then, supernatant was filtered by MICROKROS 20CM 0.65UM MPES (Repligen, and was concentrated by C06-E65U-07-S), MICROKROS 20CM 0.2 UM PES (Repligen, C02-P20U-05-S), and MICROKROS 20CM 0.05UM PS (Repligen, C02-S05U-05-S), respectively. Concentrated EVs were rinsed with EV storage buffer and collected in 1mL of EV storage buffer. Particles in the filtrate were pelleted by ultracentrifugation at 100,000g for for 90 min at 4 °C, and resuspended in EV storage buffer.[28]

### Size Exclusion Chromatography (SEC)

Size Exclusion Column was made by loading resin (G-Sep™ Agarose CL-6B, G-Bioscience) into Econo-Pac® Chromatography Columns (BIO-RAD) to bed volume 10 mL. The resin was washed with 20 mL of PBS before use. 0.5∼1 mL EV sample was loaded to the top of column and immediately 1 mL fractions were collected until desired fractions have been collected. To avoid the column running dry, PBS was added to top up the column.

### Nanoparticle Tracking Analysis (NTA)

The particle size and concentration were measured using a ZetaView® (Particle Metrix) Nanoparticle Tracking Analyzer following the manufacturer’s instruction. The following parameters were used for measurement: (Post Acquisition parameters (Min brightness: 22, Max area: 800, Min area: 10, Tracelength: 12, nm/Class: 30, and Classes/Decade: 64)) and Camera control (Sensitivity: 85, Shutter: 250, and Frame Rate: 30)). EVs were diluted in PBS between 20- and 200-fold to obtain a concentration within the recommended measurement range (0.5 × 10^5^ to 10^10^ per mL).

### Protein concentration measurement

Protein concentration was measured by Pierce™ BCA Protein Assay Kit (Fisher Scientific) by following manufacture’s protocol.

### Western Blotting

EVs were denatured at 70°C for 10 min in 1x NuPAGE LDS Sample Buffer (Thermo Fisher Scientific), separated on a 4–20% Mini-PROTEAN® TGX™ Precast Protein Gels (BioRad), and transferred to a PVDF membrane by using CAPS-based transfer buffer. The membrane with the blotted proteins was blocked with EveryBlot Blocking Buffer (BioRad) for 2 h and then incubated with a primary antibody at 4°C overnight. Following three washes with TBS with 0.1% Tween 20 (TBST), the membrane was incubated with secondary horseradish peroxidase-conjugated secondary antibody for 2h at room temperature. The membrane was again washed three times with TBST, and the protein bands were visualized by treating with SuperSignal West Pico PLUS chemiluminescent substrate (Thermo Scientific) and the image was captured by ChemiDoc Imaging System (BioRad). The following primary antibodies were used: anti-HA (Sigma Aldrich, H3663), anti-CD63 (Thermo Fisher, 10628D), and anti-ALIX (Proteintech, 12422-1-AP). The following secondary antibodies were purchased from Invitrogen: Goat anti-Mouse IgG (H+L) Highly Cross-Adsorbed Secondary Antibody, HRP (A16078) and Cell Signaling Technology: Goat anti-Rabbit IgG (H+L) Highly Cross-Adsorbed Secondary Antibody, HRP (A16110).

### DNase I Treatment of EVs

The 10 μL of eEVs were incubated at room temperature for 15 min with 1 U of DNase I (Zymo Research) and 1x DNA Digestion Buffer. The plasmid DNA was isolated from the EVs using Qiamp Miniprep kits and quantified by qPCR.

### Quantitative Real-time Polymerase Chain Reaction (qPCR)

qPCR was performed using Dream Taq DNA polymerase (ThermoFisher). Each reaction contains 200 μM dNTP, 500 nM each of forward/reverse primer, 200 nM probe (Table S1), 0.5 U DreamTaq DNA polymerase, 1x Dream Taq buffer A and 1 μL sample DNA in a total reaction volume of 10 μL using CFX96 Touch Real-Time PCR Detection System (BIO-RAD). The PCR amplification cycle was as follows: 95°C for 2 min; 40 cycles of 95°C for 20 seconds, 65°C for 30 seconds. The pDNA copy number were determined by absolute quantification using the standard curve method, and the copy number of EV encapsulated pDNA per vesicles was calculated based on NTA and qPCR results.

### Super Resolution Microscopy

Isolated EVs were analyzed with EV Profiler V2 Kit for Nanoimager (ONI) by following manufacture’s protocol. The imaging data was analyzed by CODI software (ONI).

## Result

### Characterization of eEVs particle numbers separated by TFF and centrifugation

In our initial experiment, we isolated engineered EVs using nine distinct methods to examine their size-related characteristics. Prior to isolating the EVs using TFF or centrifugation, we harvested conditioned media and subjected it to two rounds of centrifugation (600 xg for 30 min, followed by 2,000 xg for 30 min). Following this differential centrifugation, engineered EVs (eEVs) were isolated using three different TFF filters or two distinct centrifugation speeds (Fig. 1). The eEVs retained within the TFF filter were collected in an EV storage buffer[28] for improved yield. The TFF filtrate was then centrifuged at 100,000 xg for 90 min to pellet particles smaller than the molecular weight cut-off (MWCO) of the TFF. Large EVs (lEVs) were isolated by high-speed centrifugation at 12,000 xg for 90 min, while small EVs (sEVs) were isolated from the supernatant of the high-speed centrifuge at 100,000 xg for 90 min. Whole EVs were also isolated by ultracentrifugation at 100,000 xg for 90 min as a standard control commonly used in many studies. The particle yield of 50 nm TFF collected (>50) was highest among the tested populations, followed by the 200 nm collected (>200) and 650 nm filtrate (<650). There was not a significant difference between the yield of whole EVs and lEVs (Fig. 2A). The particle yield from filtrate of the 50 nm and 200 nm TFF was lower than other populations.

**Figure 1.**
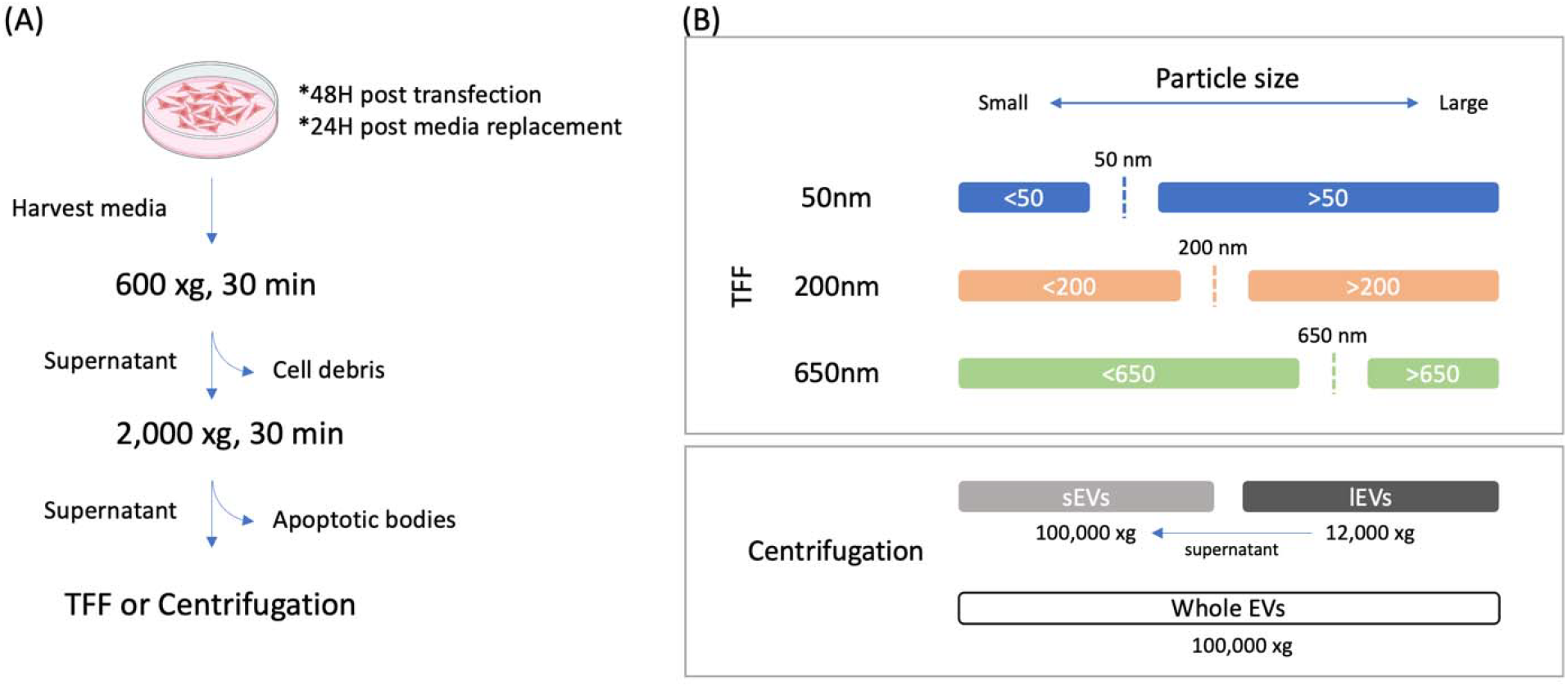
Schematic illustration of. **(A)** differential centrifugation and **(B)** EV isolation based on TFF and centrifugation.

**Figure 2.**
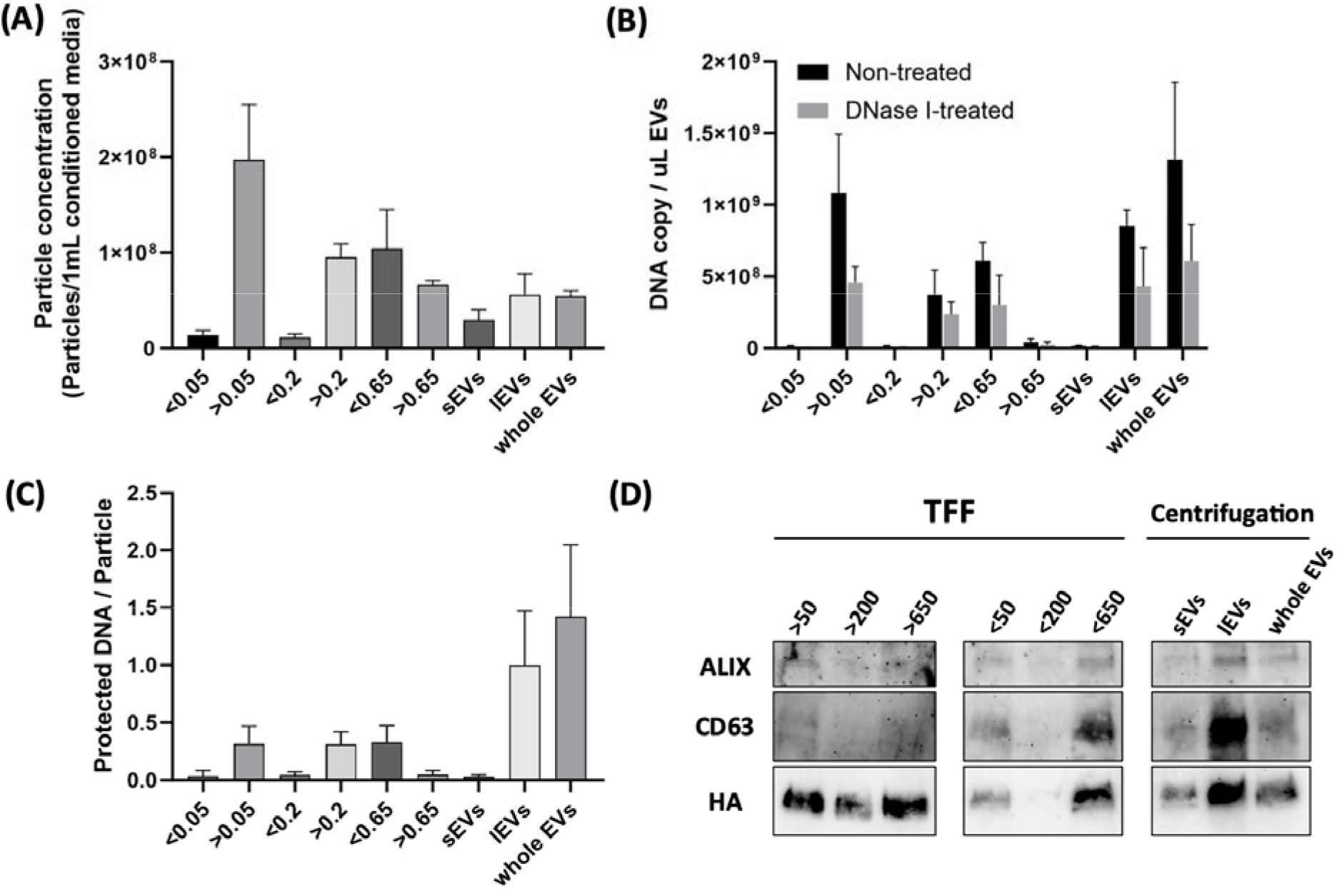
Characterization of eEVs isolated by TFF and centrifugation. **(A)** Particle concentrations per 1 mL of media isolated by each MWCO TFF or centrifugation EVs from TFF filtrate were collected by ultracentrifugation. **(B)** The DNA copy number per μL of EVs quantified by qPCR before and after DNase I treatment. **(C)** The DNA copy number per particle following DNase I treatment for each fraction. **(D)** Western blot analysis of entire particles of eEVs isolated from 8mL of conditioned media. EV marker – Alix and CD63, and engineered protein marker – HA. (A)-(C) were performed n=3. Error bars represent standard deviation.

### Quantification of plasmid DNA associated with or protected by eEVs separated by TFF and centrifugation

Subsequently, we analyzed the plasmid DNA (pDNA) associated with eEVs using qPCR. We extracted pDNA from EVs with or without DNase I treatment to quantify the total pDNA amount and encapsulated pDNA amount, respectively (Fig. 2B). The amount of pDNA per unit volume isolated by TFF was proportional to the particle yield. However, the amount of pDNA in lEV and whole EV was larger compared to TFF, even though the number of particles was smaller compared to TFF. From this data, we found that the number of pDNA packed per EV particle was approximately 1 for lEVs and whole EVs, and about 0.5 for EV groups separated by TFF, >50, >200, and <650. Almost no pDNA was detected from <50, <200, and >650, indicating that the number of pDNA per EV particle was very low (Fig. 2C).

### Protein profiling of eEV populations

To profile the isolated eEVs populations, we analyzed eEVs derived from an equivalent volume of conditioned media using Western blotting. This analysis employed standard EV markers, CD63 and Alix, as well as an eEV-specific marker, HA. Interestingly, lEV and those smaller than 650 nm (<650) exhibited higher protein levels compared to other populations. Notably, EV populations isolated by the 200 nm TFF, both smaller than 200 nm (<200) and larger than 200 nm (>200), demonstrated lower protein levels (Fig. 2D). While EV markers and the eEV marker were detected across all populations, no significant correlation was observed with the yield of particles.

### Characterization and Optimization of EV Isolation by TFF-SEC method

Our characterization and comparative analysis of eEVs populations suggested that the eEVs encapsulating pDNA exist within the 200 nm to 650 nm range, based on the MWCO of TFF filter. Interestingly, the detected amount of EV markers in eEVs separated with a 200 nm TFF was significantly lower than that with 50 nm and 650 nm (Fig. 2D). Subsequently, we evaluated a method to separate and concentrate EVs by combining 50 nm and 650 nm TFF filters, followed by SEC separation (Fig. S1). Given that protein contamination in EV isolation by TFF is reportedly lower than that by centrifugation[10], we assessed the degree of residual protein contamination by TFF and the separation capability of SEC.

Media harvested from cell culture dish underwent differential centrifugation, and the supernatant was processed using the TFF-SEC method. The filtrate was recovered using 650 nm TFF, concentrated to 1 mL using 50 nm TFF, and then recovered using PBS. After measuring the particle and protein concentrations of the collected EVs, the EVs were loaded into the SEC, and fractions were collected in 1 mL increments. Each fraction was measured using the NTA, BCA assay, and qPCR. Post-enrichment with TFF, the EVs contained approximately 0.07 ug/mL protein, but no protein contamination was detected in the fraction containing EVs recovered by SEC (Fig. S2). A slight increase in absorbance in fraction 10 on BCA assay, but it was not significant compared to the baseline that suggest the proteins observed post-TFF are eluted in fractions 10 and after.. An EV storage buffer was used during EV recovery from TFF, and EV purification was performed using SEC. The use of EV storage buffer during EVs collection increased the yield of EVs (Fig. 3, Fig. S2). The particle concentration and protein concentrations of each fraction recovered by SEC showed that the peak of EVs and the protein from the EV storage buffer were distinctly separated, albeit with slight overlap (Fig. 3A, 3B). Conversely, the peak of pDNA in each fraction overlapped perfectly with the peak of EVs, indicating that pDNA was internalized in eEVs since the DNase treatment removed free DNA prior to SEC (Fig. 3C). To confirm the effect of the centrifugation method on SEC, EVs isolated by ultracentrifugation were purified by SEC, and the particle number and pDNA content in each fraction were similarly evaluated (Fig 3D). As with EV isolation by TFF, particles and pDNA were separated into the same fraction, confirming that the EV isolation method does not affect SEC.

**Fig. 3.**
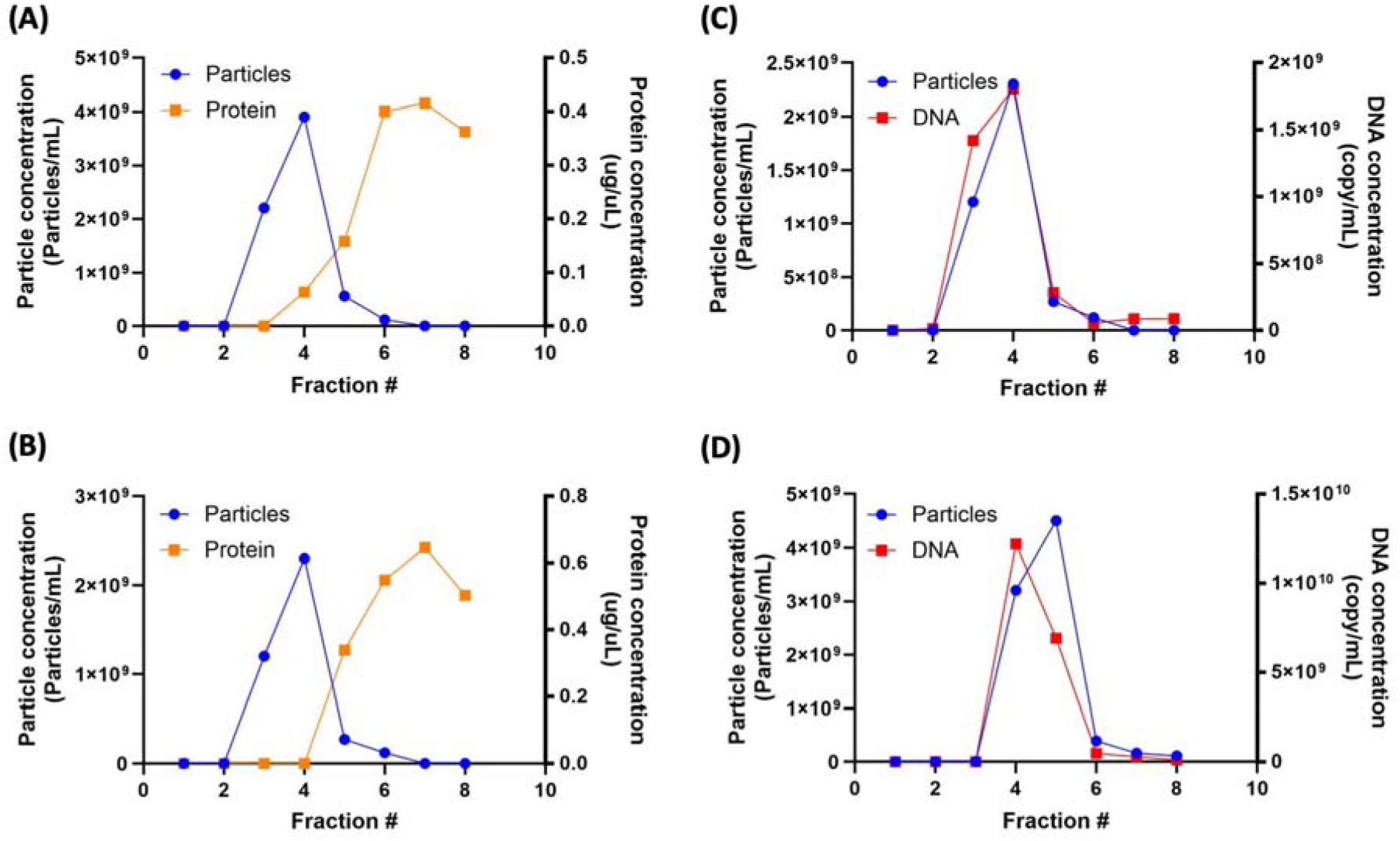
Analysis of eEVs purified further by Size Exclusion Chromatography (SEC) following TFF concentration. 1 mL of TFF (between 50 and 650 nm) isolated EVs were loaded into the SEC column. Each 1 mL fraction was collected and analyzed. **(A)** Particle concentration and Protein concentration of Naïve EVs **(B)** Particle concentration and protein concentration of engineered EVs (RDG-EVs). **(C)** Particle concentration and protected DNA amount of engineered EVs (RDG-EVs). This experiment was performed n=3, and one of the data was presented. **(D)** Particle concentration and protected DNA amount of engineered EVs (RDG-EVs) that were isolated by Ultracentrifugation and SEC.

### Single particle analysis of eEVs isolated by TFF-SEC

In our final evaluation of the method, fractions containing EVs were assessed. The fractions containing naive EVs or eEVs separated by TFF-SEC were concentrated using Amicon 100 kDa centrifugal filter and evaluated by NTA, Western blotting, and super-resolution microscopy (Nanoimager, ONI). A size distribution with a peak around 120 nm was observed for both naive EVs and eEVs (Fig. 4A, B). Traditional EV markers were detected on both Naïve and eEVs, engineered marker was detected on only eEVs (Fig S3). Co-localization of EV and eEV markers was confirmed by Super-resolution microscopy, and approximately 80% of the captured eEVs were HA positive, suggesting efficient EV surface display (Fig. 4C-F).

**Fig. 4.**
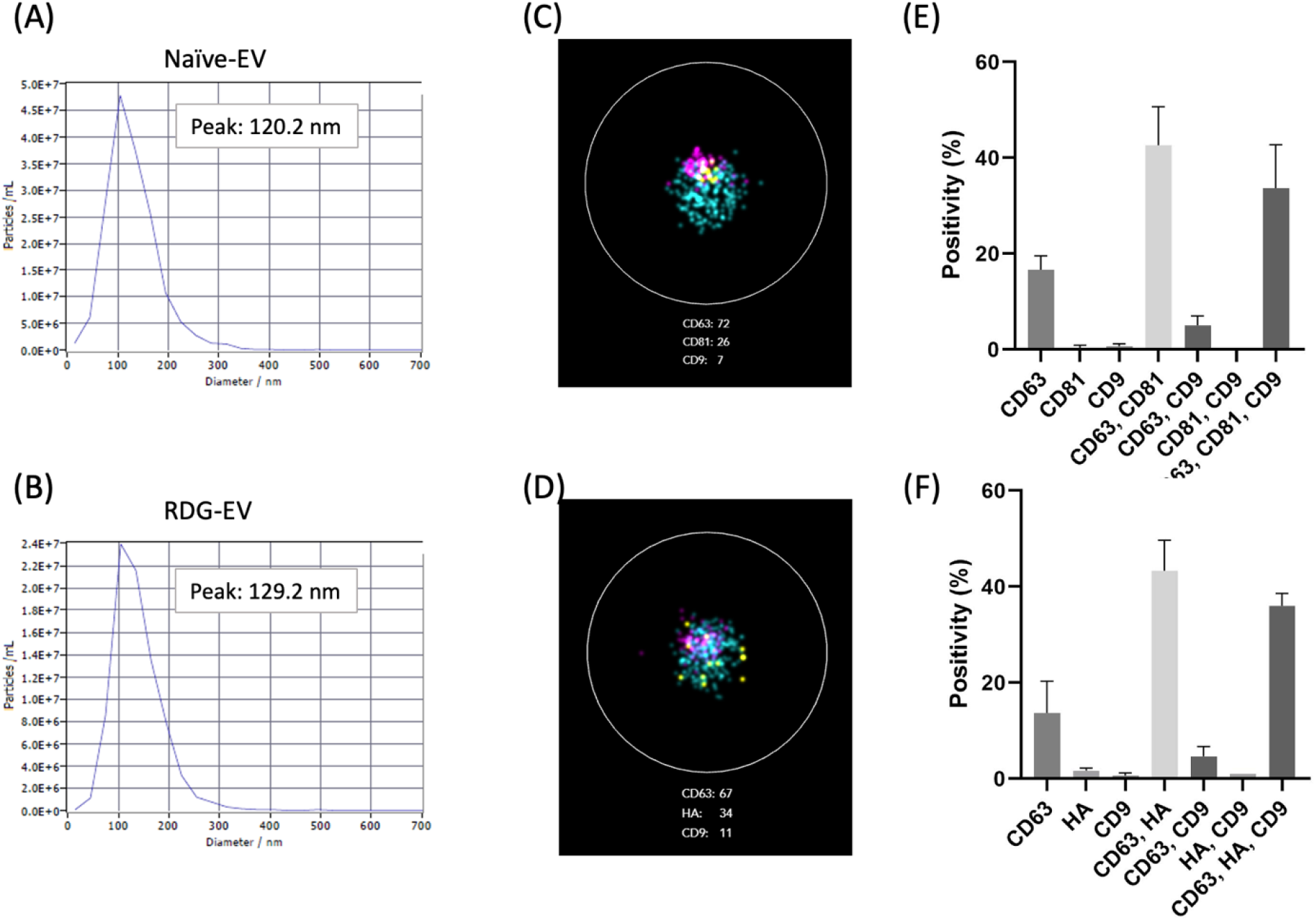
Characterization of TFF-SEC eEVs. Size distribution of TFF-SEC Naïve-EV **(A)** and RDG-EV **(B)** by NTA. Representative image of super-resolution microscopy of TFF-SEC naive-EV **(C)** and RDG-EV **(D)**. The positivity rate of EV marker (CD63, CD9) and engineered EV marker (HA) of TFF-SEC Naïve-EV **(E)** and RDG-EV **(F)**. Three individual areas on assay chip were imaged to standardize the positivity ratio (n=3). Error bars represent standard deviation

## Discussion

In the field of EV research, the efficient isolation of highly pure EVs from complex biological samples is a crucial step in various biomedical applications ranging from biomarker discovery to therapeutic development.[30-32] The current research utilizes various EV isolation methods, and existing literature underscores the widespread use of both TFF and SEC. However, many studies focus on collection of entire EV populations in the sample for analysis without delving into the detailed analysis of each EV size.[33, 34] In particular, no study has been performed on the subpopulations of EVs that contain exogenously introduced DNA cargo. We previously reported that eEVs are delivered to target sites while retaining pDNA inside, a property that could potentially enable targeted functional delivery.[29, 35] In this study, we present a comprehensive approach for large-scale eEV isolation by integrating TFF and SEC to characterize a subpopulation rich in pDNA cargo using an eEV size analysis and propose an efficient method to collect the desired EV subpopulation.

TFF has emerged as a powerful tool for the purification and concentration of biomolecules owing to its capacity to efficiently separate components based on size while minimizing sample damage.[36] TFF uses a semi-permeable membrane and a pressure gradient to effectively remove unwanted contaminants while retaining desired EVs.[37] The TFF process offers significant advantages for handling large volumes of biological samples, as evidenced by prior research in diverse fields including bioprocessing and biopharmaceutical manufacturing. [36, 38] In conjunction with TFF, SEC further enhances the isolation process by enabling the separation of EVs based on their hydrodynamic radius. In SEC, molecules are separated based on their ability to enter porous beads packed within the chromatography column. The larger molecules are excluded from entering the beads, allowing smaller molecules to pass through more easily. This differential exclusion process results in effective separation according to molecular size.[39] SEC has been extensively utilized for the purification of proteins, nucleic acids, and other biomolecules owing to its high resolution and gentle elution conditions, and is increasingly utilized for EV isolation. [40-45]

The combined use of TFF and SEC offers several advantages over traditional isolation methods. Firstly, it allows for processing large volumes of biological samples with minimal sample handling, thereby reducing the risk of sample degradation and contamination.[46] Moreover, for target biomolecular discovery or pharmaceutical-scale applications, scalability plays a pivotal role. A recent study has underscored the feasibility of large-scale EV purification through the combined utilization of TFF and size-exclusion chromatography.[24] TFF-SEC offers high resolution and gentle elution conditions, making them valuable tools for efficiently handling substantial volumes of biomolecules in laboratory and pharmaceutical settings. Furthermore, the modular and versatile nature of both techniques allows for tailoring the isolation protocol to meet the specific needs of the target EV populations and the sample matrix. For instance, TFF can be tailored to accommodate different membrane materials and pore sizes, while SEC columns can be selected based on the desired separation range and resolution. [33, 47] Such flexibility enhances the adaptability of the approach to diverse biological sources and experimental conditions. On the other hand, TFF is not suitable for applications where EVs need to be isolated from multiple or large numbers of samples, such as clinical research. SEC can be combined with commonly used ultracentrifugation methods, and, at least in this study, no significant differences in eEV characteristics were observed compared to TFF-SEC. Therefore, depending on the study design, centrifugation-based EV isolation methods may be chosen to reduce overall costs.

Our analysis showed that pDNA was mainly associated with particles above 200nm given the lack of pDNA in <200 TFF and sEV ultracentrifugation fraction. Furthermore, there was little pDNA detected in the >650nm TFF fraction. This suggested that medium sized particles isolated by lEV centrifugation or between 200nm and 650nm TFF had the largest amount of pDNA (Fig. 2A-2C). On the other hand, EVs isolated by 200 nm TFF had significantly lower amounts of protein detected by Western blot analysis (Fig. 2D), even though particles were detected by NTA. This is possibly due to physical interference between the filter and eEVs, as the filter size of 200 nm is close to the EV size peak.[48, 49]. Since EV markers and engineered markers have been detected even in populations with minimal DNA content, it suggests that subpopulations with and without pDNA exist. From these analyses, it is inferred that the pDNA-rich subpopulation has a TFF MWCO of >200 nm and <650 nm. Focusing on the yield of EVs and the ratio of stored pDNA, EVs isolated by centrifugation-based methods (lEVs, whole EVs) had a higher pDNA encapsulation rate than EVs isolated by TFF. When considering the delivery of encapsulated pDNA to cells, there is room for future research to clarify how differences in pDNA encapsulation rate affect delivery. Our results corroborate previous work, demonstrating that protein contamination in EVs obtained via TFF is lower than that from UC.[10, 24, 50, 51] However, our research extends this understanding by revealing that detectable protein contamination in EVs that is not directly bound to EVs remains. This additional insight highlights the need for continued investigation into the sources and mechanisms of protein contamination during EV isolation processes.

Notably, our approach preserves the isolated extracellular components’ structural integrity and biological activity by employing mild conditions throughout the isolation process. This ensures the isolated molecules retain their functional properties, facilitating downstream analyses such as functional assays, biomarker identification, and therapeutic development.

In a previous study, we highlighted the importance of storage buffers in EV study.[28] In this study, we found that the EV storage buffer also contributed to maintaining EV yield during the TFF step. Given that the EV fraction is eluted into PBS by buffer exchange in SEC, adding the components of the EV storage buffer before storage is desirable if the sample is not used immediately after EV collection. As mentioned earlier, we demonstrated that EVs contain pDNA. Notably, the complete overlap of EV and pDNA peaks in SEC confirms this observation from a different angle (Figure 3C). The EV fraction isolated by TFF-SEC was a homogeneous EV population with a peak at around 120 nm, predominantly comprising surface-engineered EVs.

## Conclusion

Overall, TFF-SEC represents a robust and efficient strategy for large-scale EV isolation, broadly applicable in biomedical research and biotechnology. By combining the complementary strengths of these techniques, we demonstrated the optimal conditions for engineered EV isolation that can achieve high purity, scalability, and biological activity in isolated EVs. This study and numerous prior research will pave the way for advances in EV-based disease diagnosis and drug delivery.

## Ethics approval and consent to participate

Not applicable, as no patient data was used in this research. Cell line used in this study is not relevant material under the Human Tissue Act, so no ethical approval was required.

## Consent for publication

Not applicable, as no patient data were used in this research.

## Availability of data and materials

The data that support the findings of this study are available from the corresponding author upon reasonable request.

## Authors’ contributions

Conceptualization: MH

Investigation and data generation: YH

Writing-original draft preparation: YH, VN

Writing, reviewing, and editing: YH, MH

Funding acquisition and supervision: MH

