## Supplemental for "Scalable Isolation of Surface-Engineered Extracellular Vesicles and Separation of Free Proteins via Tangential Flow Filtration and Size Exclusion Chromatography (TFF-SEC)"

Table S1. A list of primers and a probe used for qPCR

| Name | Sequence (5'-3') |
|---|---|
| qRDG_F (Forward primer) | TGTACGCTGTAACCGGTCG |
| qRDG_R (Reverse primer) | CCTGAGACGGTTTGTCGATTT |
| qRDG_P (Probe) | TCCAGCCGTCCAATCAGCATCAAT |

Fig. S1 Schematic illustration of TFF-SEC procedure

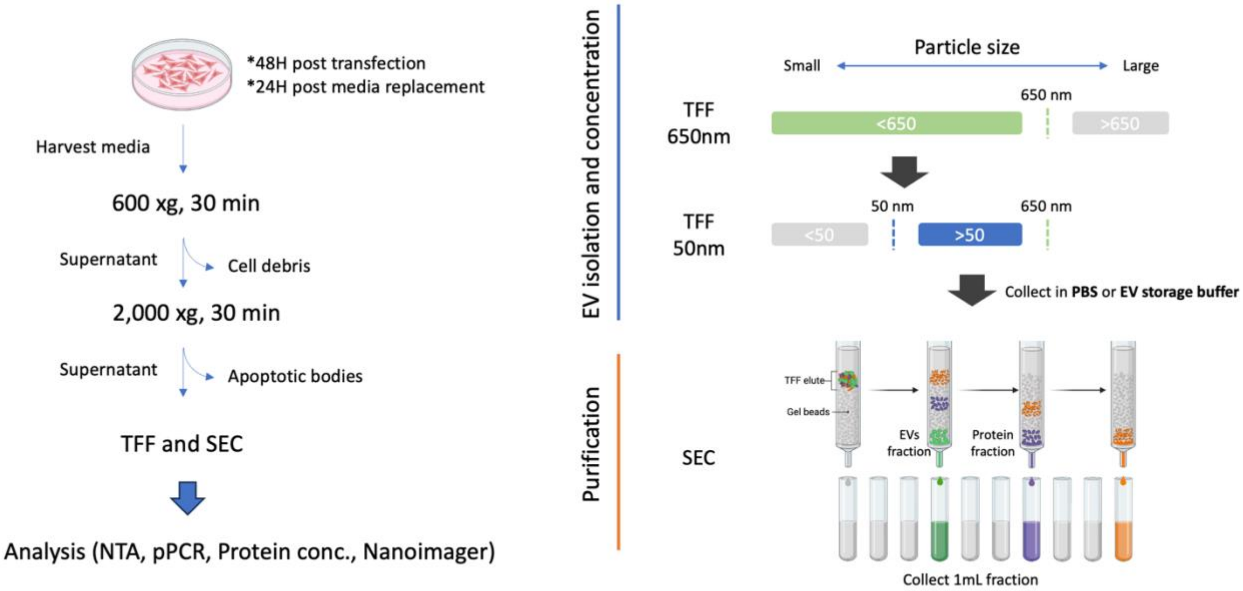

Fig. S2 SEC removed protein contamination

1 mL of TFF (between 50 and 650 nm) isolated EVs in PBS were loaded into the SEC column. Each 1 mL fraction was collected and analyzed. Particle concentration and protein concentration were measured at TFF isolation and after SEC.

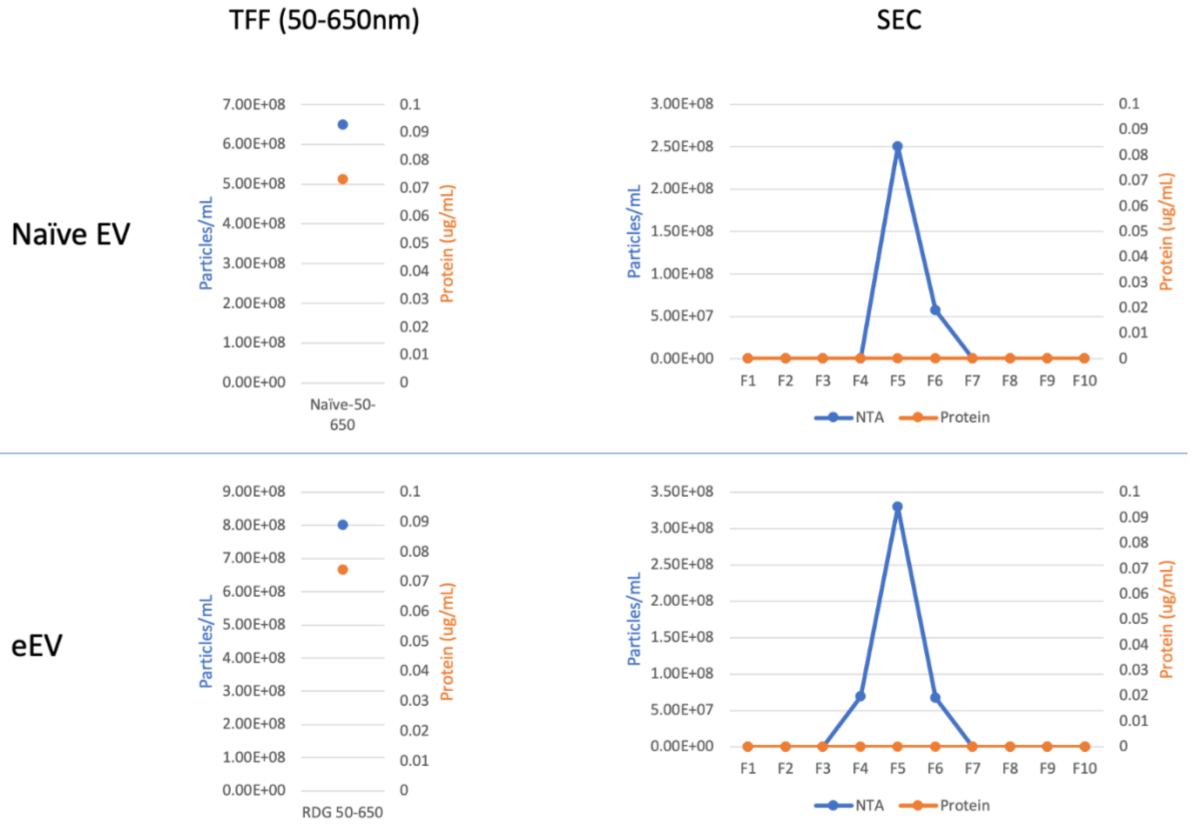

Fig. S3 Western blot analysis of TFF-SEC EVs

TFF-SEC EVs were analyzed with traditional EV markers (CD63, Alix) and Engineered EV marker (HA). Whole blot and amino black staining.

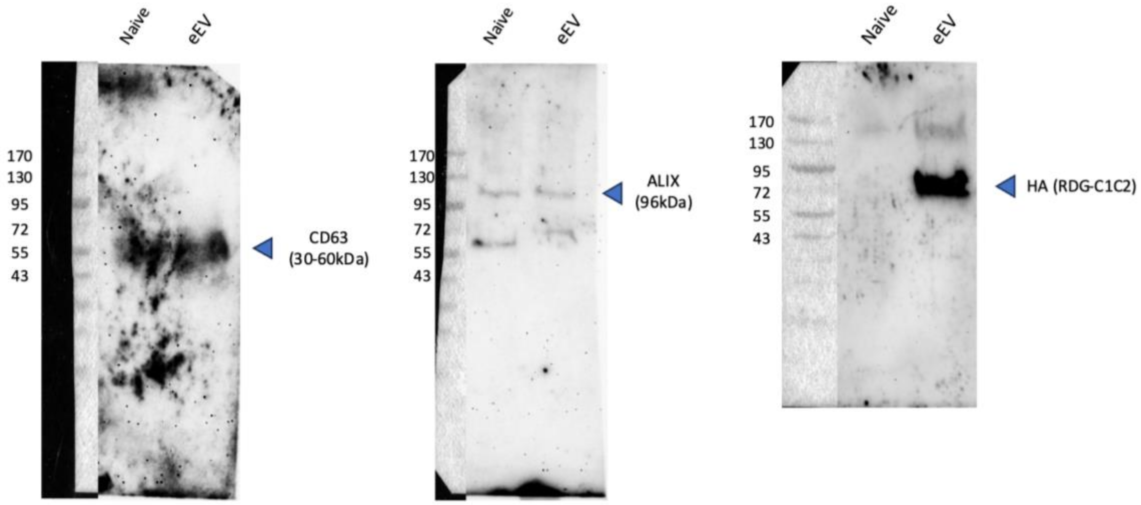

Fig. S4 whole blot of western blot analysis

Whole blot of Western blot analysis.

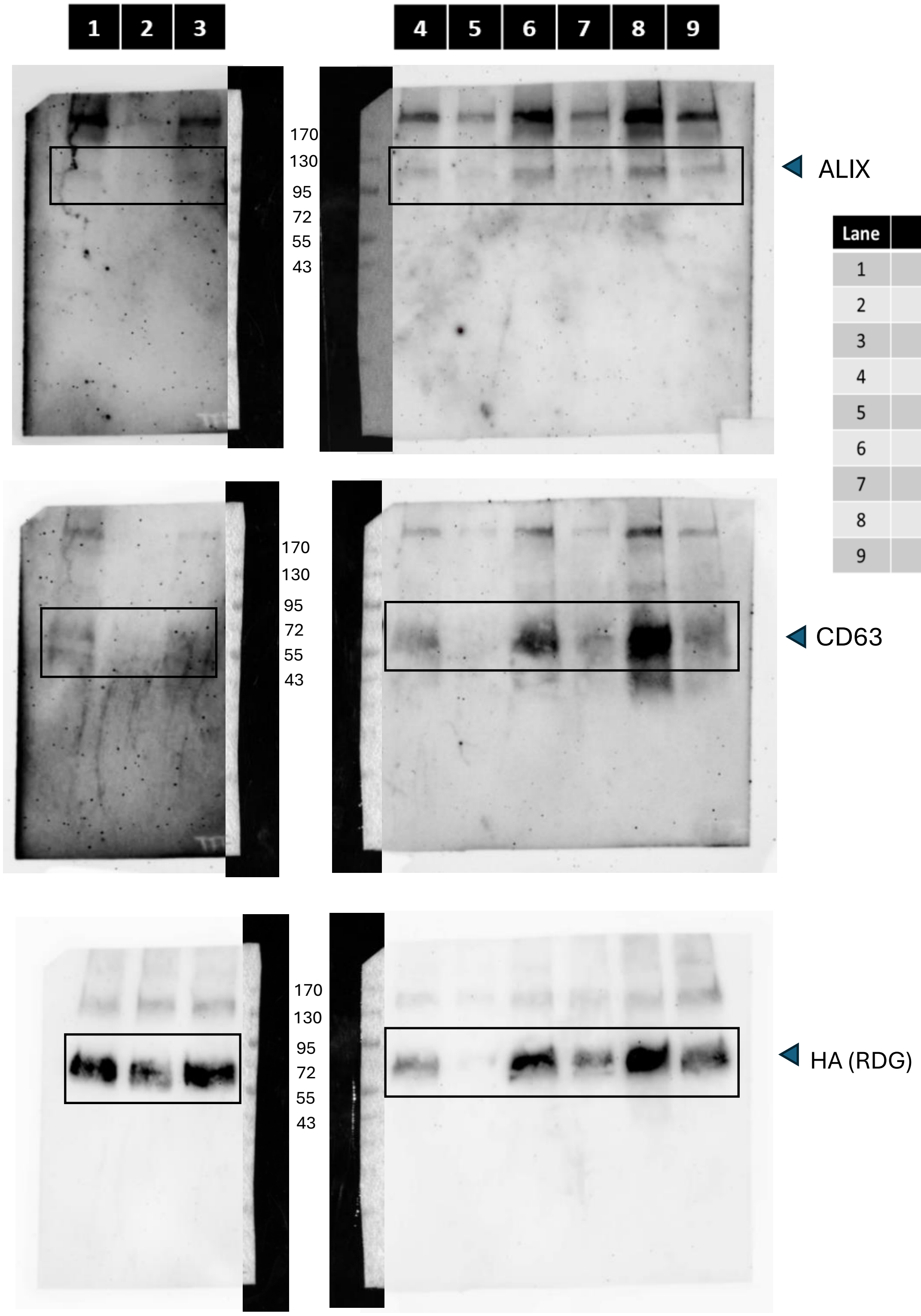


| Lane | Sample |
| --- | --- |
| 1 | TFF: >50nm |
| 2 | TFF: >200nm |
| 3 | TFF: >650nm |
| 4 | TFF: <50nm |
| 5 | TFF: <200nm |
| 6 | TFF: <650nm |
| 7 | sEVs |
| 8 | lEVs |
| 9 | Whole EVs |

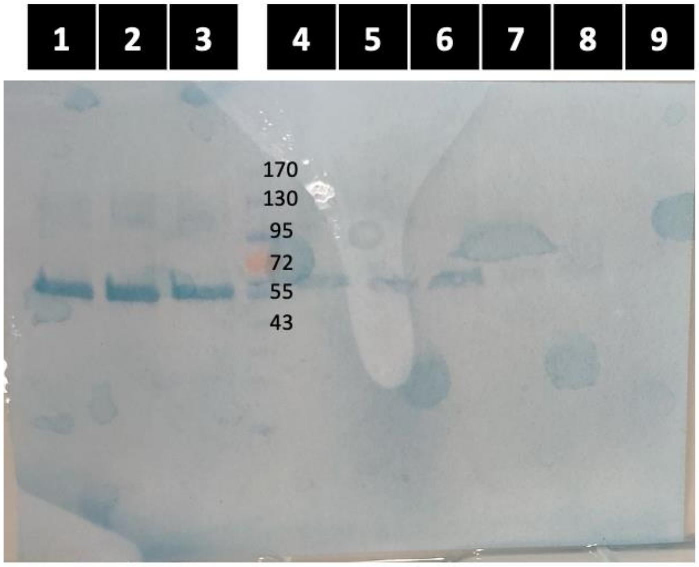
1
2
3
4
5
6
7
8
9
170
130
95
72
55
43
